# An early mTOR-dependent window during human T cell activation programs T cell state

**DOI:** 10.64898/2026.01.29.702520

**Authors:** M Valeria Lattanzio, Anouk P Jurgens, Arie J Hoogendijk, Carmen van der Zwaan, Leyma Wardak, Kaspar Bresser, Monika C Wolkers

## Abstract

T cell activation results in profound proteome remodeling that programs T cells into distinct cellular states. The mechanistic target of rapamycin (mTOR) biases T cell differentiation toward a cytotoxic fate at the expense of memory-precursor formation, making mTOR inhibition an attractive strategy to boost T cell memory during vaccination. Here, we used matched time-resolved mRNA sequencing and quantitative mass spectrometry to define how the human T cell proteome is remodeled during the first 24 hours of activation. We found that human T cells rapidly remodel their proteome in distinct, temporally ordered modules that drive translation and proliferation while promoting a cytotoxic T cell state. Notably, mTOR inhibition during the first 24 hours of T cell activation perturbed these protein modules. Strikingly, transient mTOR inhibition limited to the first 16 hours of T cell priming was sufficient to imprint a memory-like T cell state, while preserving the capacity to produce inflammatory cytokines and mediate target cell killing. Together, these findings indicate that mTOR activity dictates stable functional trajectories during early T cell activation, revealing a therapeutic window to refine vaccination responses.

## INTRODUCTION

T cells are critical players in adaptive immune responses against pathogens and malignancies. To become proficient effector cells, T cells first undergo rapid and profound changes, both during naïve T cell priming and during activation of differentiated T cell subsets. Specifically, T cell receptor (TCR) engagement triggers a cascade of signaling events that promote cell-cycle progression, induce extensive metabolic rewiring, and ultimately lead to lineage differentiation and acquisition of effector functions^1–3^.

To meet the demands of T cell expansion and differentiation, protein production is substantially increased^4–6^ and the composition of the proteome is altered in a highly dynamic fashion^7–9^. A straightforward mechanism to facilitate this enhanced protein production is to increase the availability of mRNA templates for translation. However, recent evidence indicates that this is not the primary driver for protein production. For example, activation of naïve T cells results in a ∼1.4-fold increase in total mRNA abundance but a ∼5-fold increase in total protein abundance^7^. Furthermore, the reported correlation coefficient between protein and mRNA abundance in T cells is within the range of r = 0.41-0.65^7,10,11^. These findings highlight that, in addition to transcriptional regulation, post-transcriptional events substantial contribute to proteome upon T cell priming and activation^12–15^. Despite these efforts, the precise temporal coordination of gene and protein expression during the early events of human T cell activation remains incompletely defined.

The kinase mechanistic target of rapamycin (mTOR) is a central regulator of T cell activation. Engagement of the TCR and co-stimulatory molecules trigger mTOR signaling^16,17^ leading to metabolic reprogramming^18,19^, proliferation^20^, differentiation^21–23^, survival and protein translation^24,25^. mTOR is the core component of two complexes, mTORC1 and mTORC2, which exert different functions^26,27^: Whereas mTORC1 enhances cap-dependent translation through phosphorylation of 4E-BP and LARP1^28,29^, mTORC2 regulates cellular survival and proliferation^30,31^. Coordinated mTORC1 and mTORC2 signaling promotes T cell differentiation, driving naïve CD8^+^ T cells toward short-lived effector cells^22^, while directing CD4^+^ T cells into distinct helper T cell subsets^17,23^. Because mTOR inhibition promotes the generation of predominantly memory-precursor T cells during immune responses, it has been proposed to include mTOR inhibition in vaccination strategies to enhance memory formation^21,32^. However, the precise timeline during which mTOR activity instructs cell state during T cell activation remains incompletely understood.

In this study, we measured the core transcriptional and proteomic changes in human effector T cells during the first 24 hours of TCR engagement. Time-resolved mRNA sequencing and mass spectrometry analysis revealed that proteome remodeling during T cell activation occurs in sequential phases, with tightly instructed timelines for specific gene classes. mTOR signaling specifically instructs the programs for proliferation, translation, and effector function. Importantly, transient inhibition of mTOR during T cell priming was sufficient to induce a memory-precursor-like T cell phenotype while preserving the T cell effector function. Collectively, our findings provide insights in how mTOR modulates the proteome in activated T cells, which should help guide the rational design of future vaccination strategies.

## RESULTS

### Proteome remodeling during T cell activation occurs in a staged manner

We first studied the kinetics of mRNA and protein expression during the first 24 hours of human effector T cell (Teff) activation. Teff cells were generated by activating peripheral blood-derived CD3+ T cells with anti-CD3/CD28 antibodies for three days, followed by a 9-day culture period in the absence of TCR triggering (see **Methods**). Next, Teff cells were re-activated with anti-CD3/CD28 antibodies for 2, 4, 6, or 24 hours, and subjected to mRNA-sequencing and data-independent acquisition (DIA) LC-MS/MS (**Figure 1a, Supplementary Figure 1, Supplementary Figure 2a-b**). As T cell activation results in increased cellular abundance of both mRNA and protein^7,8^, all samples obtained across the different time points were normalized, enabling comparison of relative mRNA and protein abundances.

**Figure 1.**
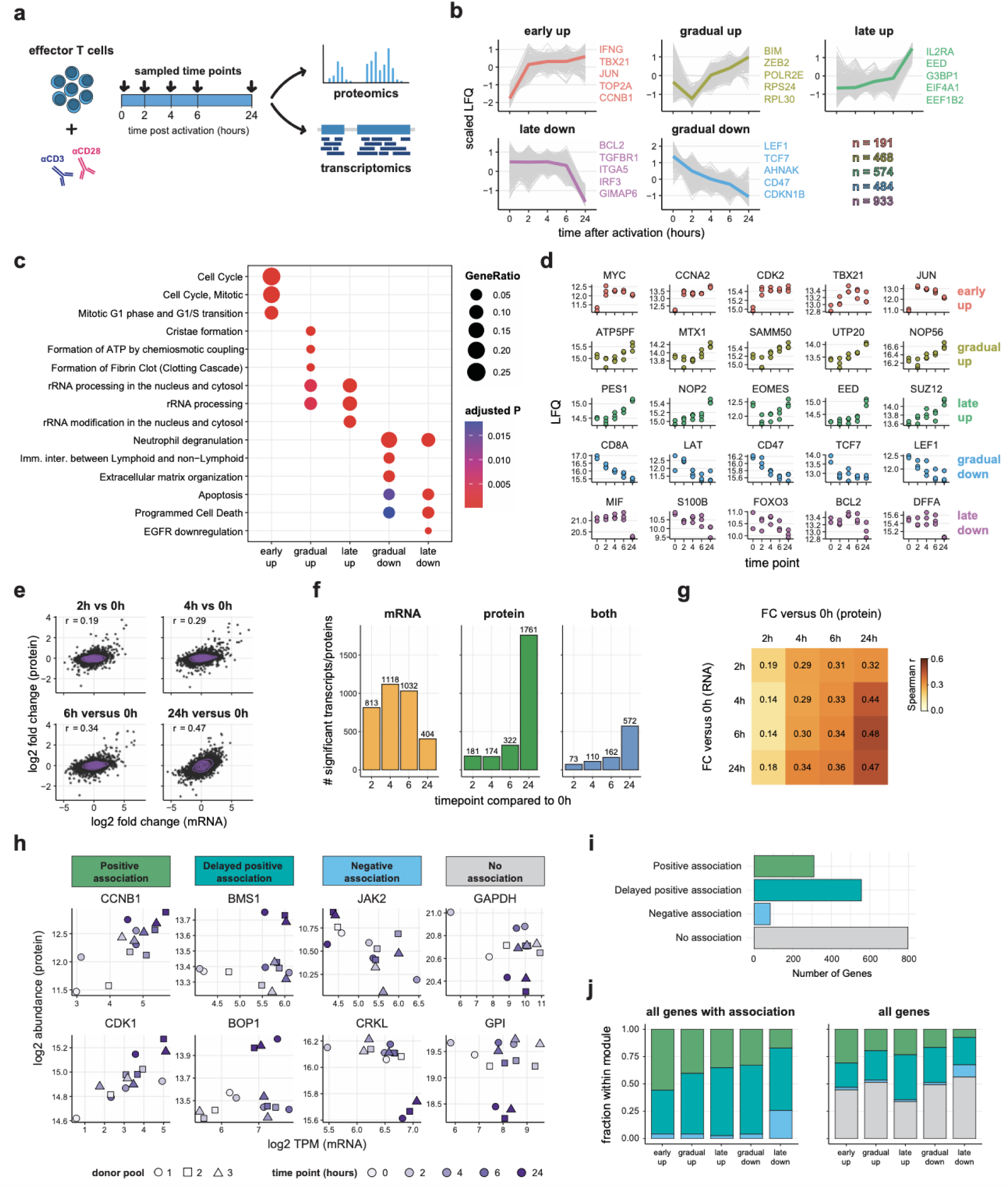
Proteome and transcriptome remodeling during early T cell activation. (**a**) Experimental design. (**b**) Differentially expressed proteins (n = 2,375, FDR < 0.05) were clustered into five temporal expression modules using hierarchical clustering. Grey lines indicate individual proteins; colored lines indicate medians. Example proteins from each module are indicated beside the plots. (**c**) Enrichment of Reactome pathways in each module. A maximum of 5 pathways is shown for each cluster. See **Supplementary Table 2** for full list. (**d**) Selected proteins for each protein module. Dots indicate individual donors. (**e**) Relationship between protein- and mRNA-level fold changes comparing each time-point to the pre-activation baseline. Dots indicate genes, colored contours indicate a 2D kernel density estimation. Spearman correlation coefficient is indicated in the plots. (**f**) Number of significant (adjusted P < 0.05) transcripts and proteins at each indicated timepoint compared to the pre-activation baseline. (**g**) Spearman correlations between the fold-changes of protein and mRNA across all activation time-points compared to the pre-activation baseline (0 hours). Correlations on the diagonal correspond to the plots shown in panel (e). (**h**) Examples scatterplots for the within-gene mRNA and protein associations. See methods for definition of associations. (**i**) Number of genes within each association. (**j**) Contribution of all positive and negative associations (left) and contribution of associations to all genes (right) indicated in (b). Displayed LC-MS data obtained from 3 pools of 40 donors.

We first asked how relative protein abundances changed during the first 24 hours of activation. To visualize this, we clustered all differentially expressed proteins (adjusted P value < 0.05; 2,375 out of 6,750 detected proteins) into 5 expression modules. Each module captured a distinct kinetic pattern (**Figure 1b, Supplementary Table 1**), including proteins that were rapidly induced (early up), proteins that progressively increased (gradual up), proteins induced only at 24 hours (late up), proteins that decreased progressively (gradual down), and proteins decreased only at 24 hours (late down). Intriguingly, these 5 protein modules were enriched for distinct functional groups of proteins (**Figure 1c-d**): (1) early up included cell cycle related proteins (e.g., MYC and CDK2) and key regulators of effector function JUN and TBX21; (2) gradual up was enriched for ribosomal RNA processing (e.g., NOP56 and UTP20) and proteins involved in ATP synthesis (e.g., MTX1 and ATP5PF); (3) late up contained proteins involved in rRNA processing and epigenetic regulation (e.g., EED and SUZ12); (4) gradual down was marked by components involved in immune-regulation and TCR-signaling components (e.g., LEF1 and LAT); and (5) late down was enriched for proteins involved in cytokine signaling (e.g., MIF and S100B) and programmed cell death (e.g., BCL2 and DFFA).

In summary, and consistent with previous reports^8,13^, T cell activation induces proteome remodeling in a highly staged process, with cell cycle components—and master regulator MYC—induced within 2 hours, followed by increased expression of the translation machinery at 6 hours, and epigenetic and transcriptional programs within 24 hours.

### Transcriptome and proteome remodeling follow distinct kinetics during activation

The abundance and availability of mRNA templates are key determinants for protein production. These determinants are defined by transcriptional and post-transcriptional events^7–9^. Therefore, we performed a correlation analysis to assess how transcriptome and proteome kinetics relate during the first 24 hours of T cell activation. Interestingly, changes in mRNA and protein abundance correlated only mildly at early timepoints (0-6h, spearman r ≈ 0.27) but gradually became more aligned at later timepoints (reaching spearman r = 0.47 at 24h; **Figure 1e**). This increased alignment could, in part, be explained by delayed protein expression, as early changes in mRNA abundance showed a higher correlation with protein abundance changes at later timepoints (**Figure 1f-g**). In line with this observation, the gene set ‘ribosomal RNA processing’, which was enriched in the ‘gradual up’ and ‘late up’ protein expression modules, already peaked at 4 hours post-activation at the mRNA level (**Supplementary Figure 2c-d**).

To better understand the relationship between mRNA and protein abundance, we calculated Spearman’s rank correlation coefficients (ρ) between mRNA and protein levels for genes within each of the five protein expression modules (Figure 1b), classifying genes as positively associated (ρ > 0.5), inversely associated (ρ < −0.5), or not associated (remaining genes; **Figure 1h**). In addition, a time-lagged Spearman correlation (i.e., correlating mRNA abundance at earlier timepoints with protein abundance at later timepoints; **see Methods**) was used to account for delayed protein expression (**Figure 1h-I, Supplementary Table 3**). The majority of the positively associated genes belonged to the ‘early up’ protein module (**Figure 1j**) and were strongly enriched for several pathways involved in the cell cycle (**Supplementary Figure 3**), indicating that a tight relationship between transcription and translation contributes to these early kinetics. Genes with a delayed positive association primarily belonged to protein modules with gradual kinetics (e.g., the translation regulators *BSM1* and BOP1; **Figure 1h-i**). Almost all genes that were negatively associated belonged to the ‘late down’ protein module (e.g., cytokine signal transducers *CRKL* and *JAK1*; **Figure 1h-i**). Interestingly, the largest group of genes (1,147 out of 2,375) did not display an obvious association between RNA and protein abundance (e.g. metabolic enzymes *GAPDH* and *GPI*; **Figure 1j**). In conclusion, the relative RNA and protein abundances follow distinct kinetics upon TCR triggering, pointing to the pivotal role of post-transcriptional mechanisms during these early stages of T cell state wiring.

### mTOR instructs proteome remodeling during early T cell activation

mTOR is involved in several (post-)translational regulatory mechanisms, including the promotion of translation^11,33^, and mTOR has been shown to boost translation during early T cell activation^7,34^. Therefore, we asked if mTOR activity is involved in the kinetics of the identified protein expression modules. To this end, Teff cells (matched donor pools to Figure 1) were re-activated with anti-CD3/CD28 antibodies for 0, 6, and 24 hours in the presence of the mTORC1/2 inhibitor Torin-1 or DMSO alone. Non-stimulated T cells (0h) were incubated with the inhibitor or DMSO alone for 6 hours (**Figure 2a, Supplementary Figure 4a-e**). Torin-1 treatment during T cell activation efficiently suppressed phosphorylation of the mTOR targets 4E-BP (mTORC1) and AKT (mTORC2; **Supplementary Figure 4f-g**), confirming that mTOR activity was perturbed.

**Figure 2.**
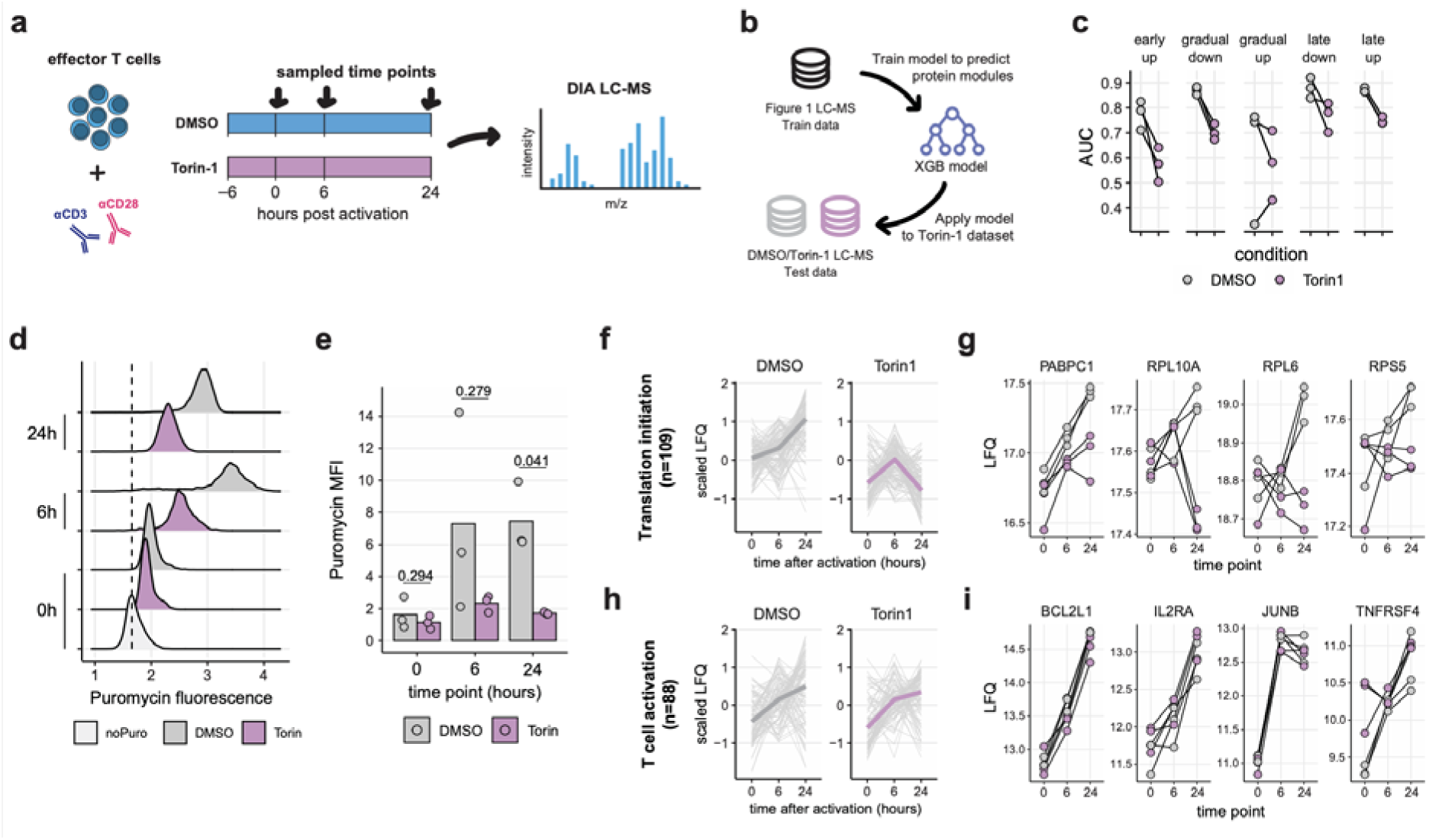
mTOR instructs proteome remodeling during early T cell activation. (**a**) Experimental design. (**b**) LC-MS data from Figure 1 was used to train a XGBoost classifier for the protein expression modules. This model was then used to classify the DMSO/Torin-1 LC-MS data. (**c**) Model performance plotted as area under the ROC curves (AUC). Black lines connect matched donor pools. (**d-e**) Puromycin incorporation assay of DMSO and Torin-1 treated CD8^+^ T cells at indicated time-points after CD3/CD28 cross-linking. Histograms display puromycin incorporation in a representative donor (d), barplots display the median fluorescence intensity for all donors (e). Black dashed line indicates the maximum density of fluorescence for the no puromycin control. (**f-g**) Expression kinetics of proteins from the Reactome pathway ‘translation initiation’ (f), and selected examples (g). Grey lines indicate individual proteins; colored lines indicate medians (f). Black lines connect matched donor pools (g). (**h-i**) Expression kinetic of proteins from the Reactome pathway ‘T cell activation’ (h), and selected examples (i). Grey lines indicate individual proteins; colored lines indicate medians (h). Black lines connect matched donor pools (i). LC-MS data is obtained from 3 pools of 40 donors. Panel (d) and (e) display 3 pools of 5 donors and are representative of 2 independent experiments. P values indicated in (e) were calculated using a two-tailed paired Student’s t-test followed by Benjamini-Hochberg correction.

We first studied which of the protein expression modules (as defined in Figure 1b) were specifically sensitive to mTOR blockade. Because the relationship between protein abundance and time may not be well described by simple statistical models, we applied a tree-based machine learning approach. In short, we trained an XGBoost model^35^ on the control dataset (Figure 1) to learn the characteristic kinetic protein expression profile of each of the 5 defined modules and used it to classify the module identity for each protein in the Torin-1/DMSO-treated samples (**Figure 2b, Supplementary Figure 5a**). This analysis revealed that Torin-1 treatment strongly affected 4 out of 5 protein modules (**Figure 2c, Supplementary Figure 5b**), indicating that mTOR signaling broadly regulates protein expression kinetics during the first 24 hours of T cell activation. In line with that finding, Torin-1 treatment globally reduced protein translation in activated T cells, as measured by the puromycin incorporation assay^36^ (**Figure 2d-e**). Moreover, LC-MS analysis revealed that the abundance of proteins related to translation initiation was strongly reduced in Torin-1 treated Teff cells (**Figure 2f-g**). Despite this general perturbation of translation, Torin-1 treatment did not block T cell activation *per se*. In fact, the abundance of proteins that are induced downstream of TCR signaling, including BCL2L1, CDK4, IL2RA, JUNB, and TNFRSF4, was upregulated normally (**Figure 2h-i**). Thus, although mTOR regulates early proteome remodeling across all the protein expression modules, its regulation is concentrated on a distinct set of proteins.

### mTOR inhibition disrupts distinct pathways

We next investigated if specific pathways were affected by mTOR inhibition, by clustering the Torin-1 sensitive proteins across all timepoints (**Figure 3a, Supplementary Table 4**) and assessing these clusters for enriched pathways (**Figure 3b-c**). Torin-1 treatment strongly repressed the induction of proteins involved in protein translation (RPS15, EEF1G; cluster C1) and to a lesser extent cell cycle (CDK4, CCNB1; cluster C2), demonstrating that these known downstream effects of mTOR inhibition can already be observed during the first 24 hours of TCR triggering. Torin-1 treatment also blocked the repression of a large group of proteins (cluster C3). These proteins are associated with several pathways, including components of cytokine signaling (IFIT5, JAK1), neutrophil degranulation (CSTB, ADAM10), and transcriptional regulation (TCF1, LEF1; **Figure 3a-c**). Finally, Torin-1 treatment induced the expression of several proteins of which the expression was stable in the DMSO control during the first 24 hours of activation (cluster C4, **Figure 3a**). No pathway enrichment was detected in cluster C4. Nevertheless, this cluster included key regulators of T cell function, including the transcription factors ARID5A and GATA3, the catalytic subunit of PI3K (PIK3CG), activator of the NF-κB pathway IKK-β (IKBKB), and cytokine signaling modulator CBLB (**Figure 3c**).

**Figure 3.**
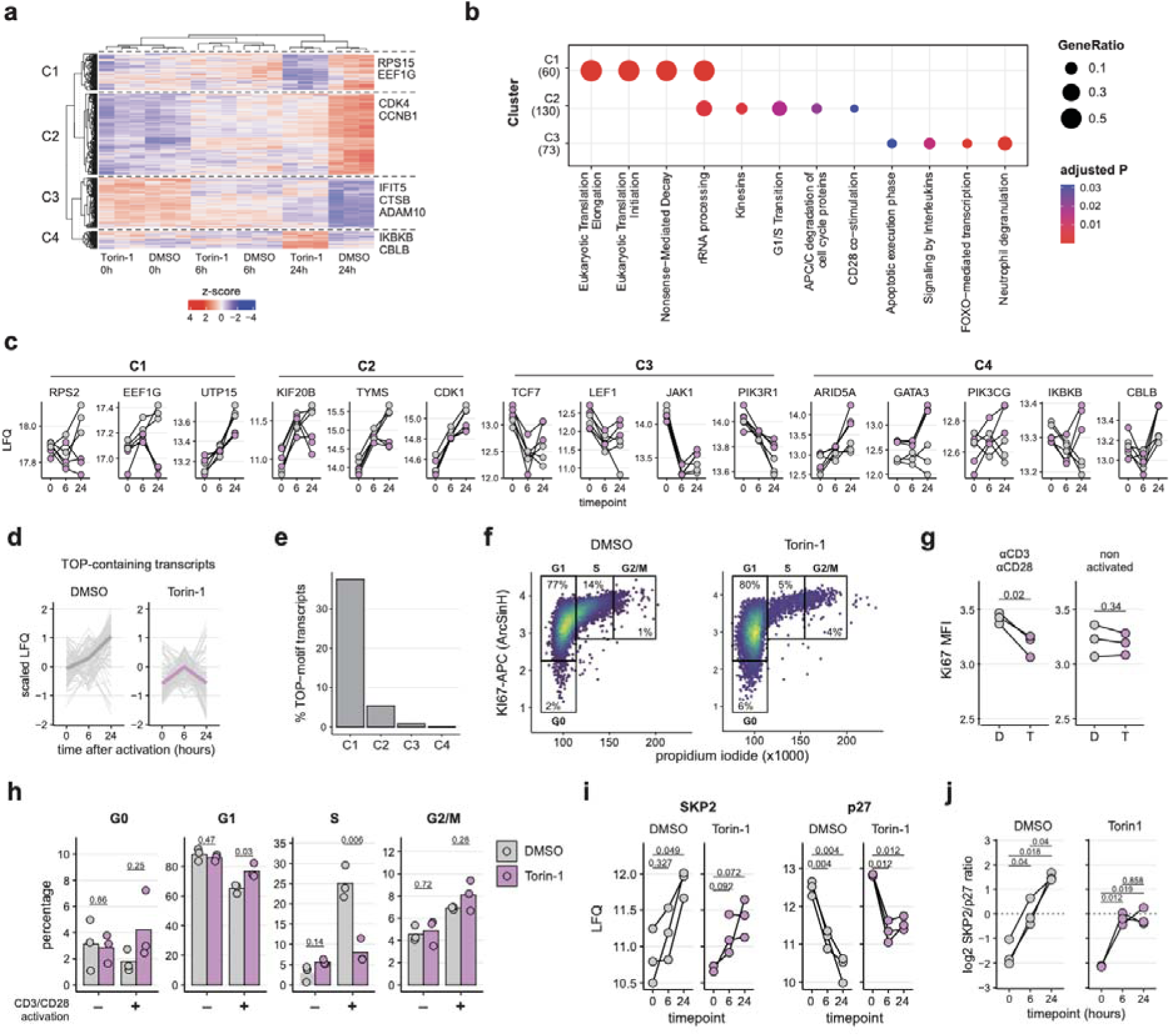
mTOR instructs early activation-induced T cell differentiation. (**a**) Clustering analysis of all Torin-1 dependent differentially expressed proteins from the LC-MS data. Euclidean distance is used for the rows; spearman correlation is used for the columns. (**b**) Enrichment analysis for Reactome pathways in each cluster using a hypergeometric model. A maximum of 5 pathways is shown for each cluster. See **Supplementary Table 5** for full list. (**c**) Representative proteins from each Torin-1 sensitive cluster. Black lines connect matched donor pools. (**d**) Expression kinetics of proteins translated from TOP-motif containing transcripts. Grey lines indicate individual proteins; colored lines indicate medians. (**e**) Percentage of TOP-motif containing transcripts within each Torin-1 dependent cluster (panel a). (**f-g**) Propidium-iodide-based cell cycle analysis of CD8^+^ T cells at 24 hours post-activation (anti-CD3/CD28). (**f**) Representative scatterplots with 2D kernel density estimation overlay to illustrate gating strategy. Boxes indicate definition of cell cycle phases. (**g**) Median fluorescence intensity of Ki67 staining. Black lines connect individual donors. (**h**) Percentage of cells in each phase of the cell cycle. Dots indicate individual donors; bars represent group means. (**i-j**) Expression kinetics (LC-MS) of SKP2 and p27 (i), and relative SKP2/p27 abundance ratios (j). Black lines connect matched donor pools. LC-MS data i obtained from 3 pools of 40 donors. Panel (f-h) display 3 pools of 5 donors and are representative of 2 independent experiments. P values indicated in (g-j) were calculated using a two-tailed paired Student’s t-test followed by Benjamini-Hochberg correction.

Next, we assessed how mTOR instructs these Torin-1 sensitive protein clusters during T cell activation. Ribosome biogenesis and protein translation pathways were strongly repressed during mTOR inhibition (cluster C1, **Figure 3b**). Furthermore, mTOR is known to promote the translation of transcripts that contain 5’UTR TOP-motifs, which is mediated through phosphorylation of the translation repressors 4E-BP and LARP^25,29^. Consistent with our observation that Torin-1 treatment effectively reduces p4E-BP (**Supplementary Figure 5**), the protein output of TOP-motif transcripts was perturbed (**Figure 3d**). Moreover, TOP-motif transcripts were strongly enriched in cluster C1 (**Figure 3e**), highlighting this pathway as a mechanism driving the Torin-1 induced repression of proteins in cluster C1.

Perturbation of mTOR signaling also affected the abundance of proteins involved in proliferation (C2, **Figure 3b**). Consequently, cell cycle progression was affected in activated T cells treated with Torin-1, as evidenced by the reduced expression levels of Ki67 (**Figure 3f-g**) and a decreased percentage of activated T cells progressing through the S-phase (**Figure 3h**). In cancer cells, mTORC1-mediated phosphorylation stabilizes SKP2, an E3 ubiquitin ligase that promotes proliferation by degrading the cell cycle inhibitor p27^37,38^. Consistent with this mechanism, SKP2 protein abundance increased upon activation of human T cells, coinciding with reduced p27 abundance (**Figure 3i**). Importantly, this alteration in the abundance ratio between SKP2 and p27 was reduced upon Torin-1 treatment (**Figure 3i-j**). These data suggest that mTOR signaling contributes to the proliferation program in activated Teff cells, at least in part, by regulating the abundance of SKP2 protein.

### mTOR instructs early activation-induced T cell differentiation

Upon activation, T cells differentiate into highly active cytotoxic or long-lived memory-precursor cells^1,3,39^. This process is coordinated by the transcription factors TCF1 (quiescence/multipotency) and T-bet (effector function). Long-term treatment with rapamycin (>7 days) or T-cell-specific ablation of mTOR complex components in murine infection skews T cells toward the TCF1^HIGH^ memory-precursor cell state^21,22,40^. In human Teff, the abundance of proteins involved in T cell differentiation was altered upon Torin-1 treatment (C3; **Figure 3a-b**). Intriguingly, flow cytometric analysis showed that a 24-hour treatment with Torin-1 during re-activation of Teff cells was sufficient to retain high levels of TCF1 (**Figure 4a-b**). Concordantly, Torin-1 treatment substantially blocked the induction of T-bet, a major driver of the effector T cell program (**Figure 4c-d**). In addition, gene-set enrichment analysis of the LC-MS data showed that proteins associated with various T cell memory subsets were enriched in Torin-1 treated samples (**Figure 4e**). This included increased expression of several hallmark proteins associated with T cell memory, such as BCL2, SELL, GATA3 and CD27, and downregulation of the effector T cell related proteins EOMES, ZEB2, and EZH2 (**Figure 4f**).

**Figure 4.**
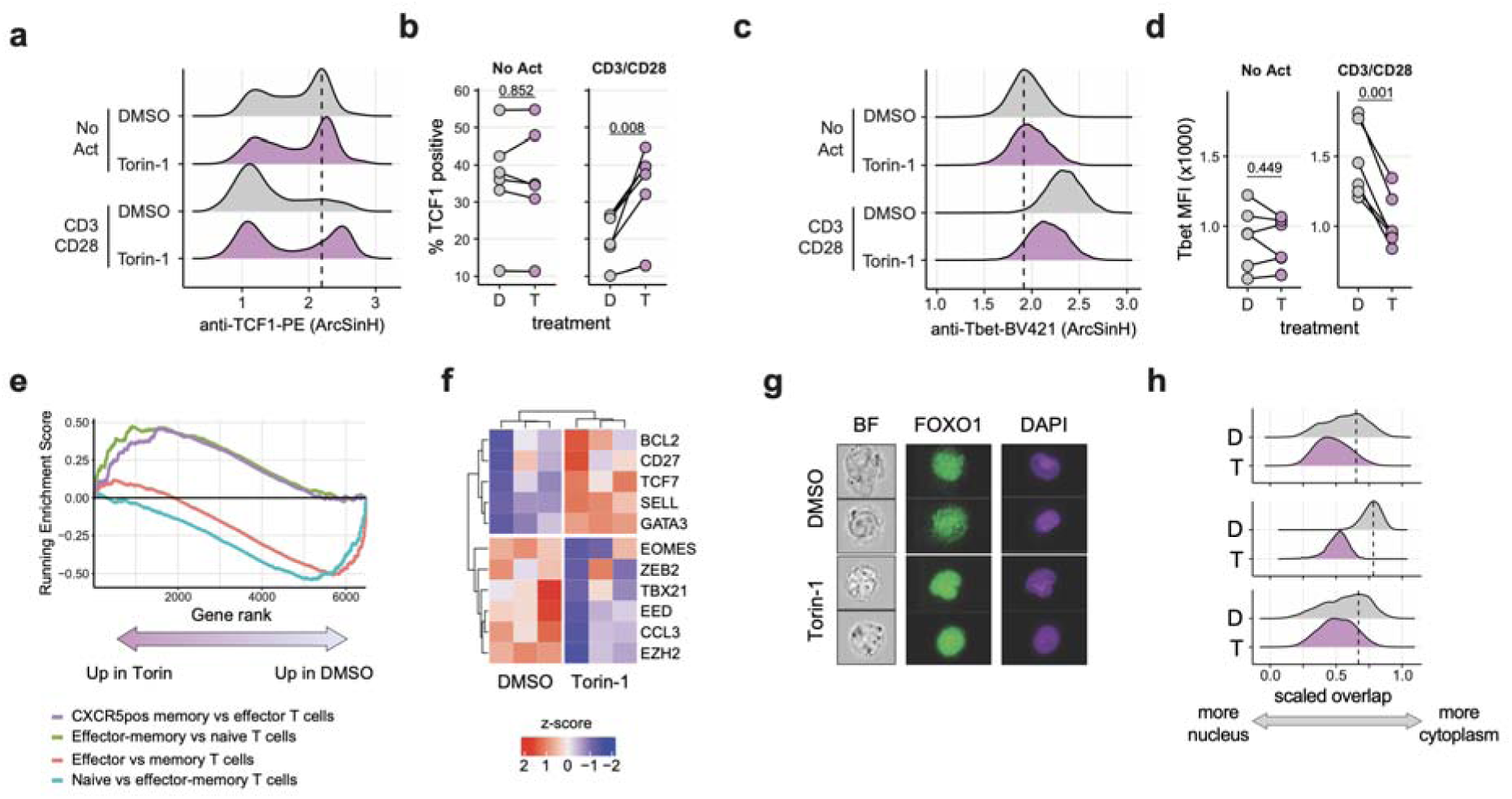
mTOR instructs early activation-induced T cell differentiation. (**a-d**) Intracellular staining for TCF1 (a-b) and TBET (c-d) protein expression of CD8^+^ T cells at 24 hours post-activation (anti-CD3/CD28), measured by flow-cytometry. Representative histograms for single donors are shown (a, c). Dashed lines indicate the maximum density of fluorescence for the non-activated control (DMSO). Strip charts display percentage of TCF1 positive cells (b) or median fluorescence intensity of TBET (d). Black lines connect individual donors. (**e**) Gene-set enrichment analysis comparing Torin-1 and DMSO treated samples. Running enrichment score of selected pathways is shown, see **Supplementary Table 6** for full results. (**f**) Clustering analysis of selected genes representative of T cell differentiation states. (**g-h**) Intracellular localization analysis of FOXO1 and nucleic acid dye DAPI of CD8^+^ T cells at 24 hours post-activation. Representative images (g) and quantifications for 3 donors (h) are shown. Overlap between DAPI and FOXO1 signals is scaled between 0 and 1 to visualize the amount of FOXO1 outside of the nucleus. LC-MS data is obtained from 3 pools of 40 donors. Panel (b, d) display 3 pools of 5 donors and are representative of 2 independent experiments. P values indicated in (b, d) were calculated using a two-tailed paired Student’s t-test followed by Benjamini-Hochberg correction.

We next studied how mTOR inhibition favors the multipotency/memory trajectory during Teff activation. Previous work in mice showed that the transcription factor FOXO1 controls the expression of TCF1. FOXO1 is directly phosphorylated through the mTOR-AKT axis^22,31^, which excludes FOXO1 from the nucleus and prohibits its transcriptional activity^41^. In line with these prior studies, Torin-1 treatment of human Teff limited AKT phosphorylation upon T cell activation (**Supplementary Figure 4**). Importantly, Torin-1 treatment also resulted in increased nuclear accumulation of FOXO1 after 24 hours of TCR stimulation compared to DMSO controls (**Figure 4g-h**). Combined, these data imply that the induction of a more multipotent T cell state by Torin-1 during Teff activation is driven by sustained nuclear retention of FOXO1 through the mTOR-AKT axis.

### mTOR inhibition during priming imprints T cell state

Having established that Torin-1 treatment during T cell activation maintains Teff cells in a more multipotent state, we next asked if mTOR inhibition would yield similar effects during naïve T cell priming, and whether this multipotent state remains ‘imprinted’. Therefore, we activated naïve CD8^+^ T cells for 48 hours using anti-CD3/CD28 beads in the presence or absence of Torin-1 (**Figure 5a**). After 48 hours, T cells were removed from activation, washed to eliminate Torin-1 from the cultures, and subsequently maintained under identical culture conditions (**Figure 5a**). At day 7 post activation, Torin-1 treated T cells had expanded significantly less than control treated T cells (5-fold versus 30-fold expansion; **Figure 5b**). Consistent with our observations in Teff cells (**Figure 4**), Torin-1 treatment during T cell priming resulted in a substantially higher abundance of TCF1^HIGH^ T cells (**Figure 5c**). Moreover, mTOR inhibition substantially increased the expression of the lymph node homing receptor CCR7 and the co-stimulatory protein CD28 (**Figure 5d-e**), whereas expression of the inhibitory receptors CD39 and TIM3 was markedly reduced (**Figure 5d-f**). Memory-precursor-like T cells generally produce less effector molecules than their more differentiated counterparts (e.g., short-lived effectors)^42^. Strikingly, Torin-1 treated T cells displayed only a slight reduction in the production of TNF after 3 hours of re-activation, and no significant differences in the production of IFNγ, or IL2 (**Figure 5g**). Together, these data indicate that Torin-1 treatment during T cell priming skews T cells towards a more multipotent T cell state.

**Figure 5.**
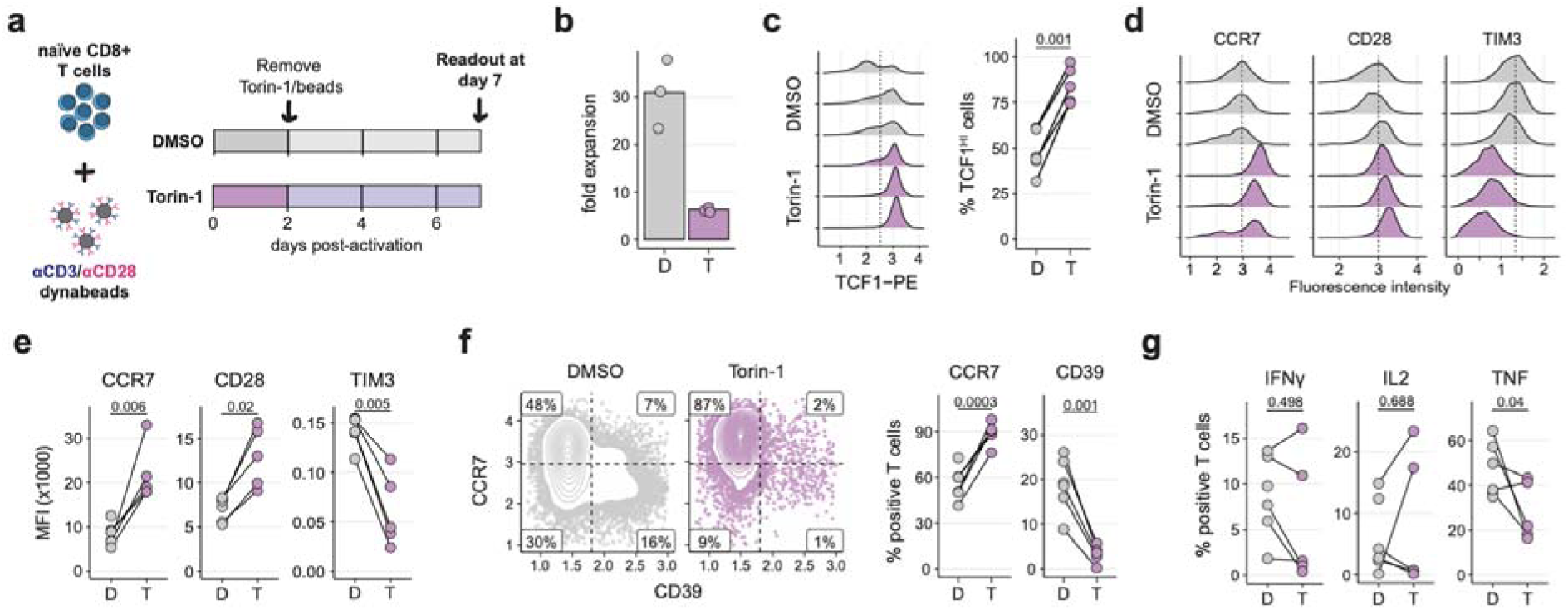
mTOR inhibition during T cell priming imprints T cell state. (**a**) Experimental setup. Naïve CD8^+^ T cells were activated in the presence of Torin-1 or DMSO for 48 hours and subsequently cultured for 7 days in the absence of both. Read-outs are performed at day 7 post-activation. (**b**) Fold-expansion of T cell cultures between day 0-7. (**c**) TCF1 protein expression, measured by intracellular flow-cytometr at day 7 post-activation. Representative histograms of TCF1 fluorescence intensity (left) and percentage of TCF1-positive T cells (right) are shown. Dashed line indicates the threshold for positive cells. (**d-e**) Surface staining for indicated proteins, measured by flow-cytometry. Representative histograms (d) and median fluorescence intensity (MFI) of indicated proteins (e). Dashed lines (d) indicate the maximum density of fluorescence for the DMSO controls, solid lines (e) connect individual donors. (**f**) Surface staining for indicated proteins, measured by flow-cytometry. Scatterplots with 2D kernel density contour for representative samples (left) and summarized percentages (right) are shown. (**g**) Intracellular staining for indicated cytokines after a 3-hour re-stimulation within anti-CD3/CD28 antibodies in the presence of brefeldin A. Solid lines connect individual donors. Displayed data is compiled from of 2 independent experiments comprising 5-6 donors. P values indicated in (c, e-g) were calculated using a two-tailed paired Student’s t-test followed by Benjamini-Hochberg correction.

### Transient mTOR inhibition during priming is sufficient to imprint a multipotent T cell state

Treatment with mTOR inhibitors has been shown to skew T cell responses to a multipotent memory-precursor fate in mouse models^21,22^. In line with this, recent clinical studies revealed that patients receiving continuous mTOR inhibition generated enhanced vaccine-induced T cell memory pools^43–45^. However, long-term treatment with immune-suppressive mTOR inhibition can lead to adverse effects^46,47^, impeding the broad use of mTOR inhibitors to improve vaccine efficacy. Because we found that a 48-hour Torin-1 treatment imprints a multipotent state in primed naïve T cells, we questioned whether transient mTOR inhibition during the initial stage of T cell priming would be sufficient achieve a similar outcome. To test this, naïve T cells were primed with anti-CD3/CD28 beads in the presence or absence of Torin-1 for 16 hours (**Figure 6a**). T cells were then washed to remove Torin-1, a and T cell activation was continued for the remaining 32 hours (total 48 hours). This transient Torin-1 treatment impacted the population expansion less than the continuous treatment (10-fold versus 5-fold expansion; **Figure 6b**). Remarkably, transient Torin-1 treatment was sufficient to imprint a more multipotent T cell state, as T cells expressed significantly higher levels of TCF1 at day 7 post-activation (**Figure 6c**). This was matched by enhanced expression of CCR7 and CD28 and reduced expression of CD39 and TIM3 (**Figure 6d-e**). Importantly, the transient Torin-1 treatment did not affect the production of cytokines during a 3-hour re-stimulation assay (**Figure 6f-g**). Yet, expression of the cytolytic protein granzyme B was markedly reduced (**Figure 6h**), which is a feature of multipotent T cells^48^. To test whether the reduced Granzyme B expression resulted in reduced T cell functionality, we measured the killing capacity of Torin-1 and control -treated T cells in a co-culture assay. To test this independent of TCR-specificity we used ‘universal target’ cells that express a membrane-tethered single-domain anti-CD3 antibody (**Figure 6i, see Methods**). Whereas the production of TNF was unaltered, T cells produced slightly less IFNγ and IL2 after 16 hours of co-culture (**Figure 6j**). Interestingly, T cell activation markers (CD25, CD137, CD69) were induced to a similar degree in T cells primed with or without Torin-1 (**Figure 6k**), suggesting that the response to TCR triggering was equal. In line with this observation, Torin-1 treated T cells were equally capable of killing target cells as the control T cells, shown by equal numbers of remaining Mel888 cells after a 48-hour co-culture (**Figure 6l**). In conclusion, transient Torin-1 treatment during the first 16 hours of T cell priming is sufficient to imprint T cell differentiation toward more multipotent T cell state. Importantly, despite lacking many effector-associated characteristics, Torin-1 treated treatment generates functionally competent, cytotoxic T cells.

**Figure 6.**
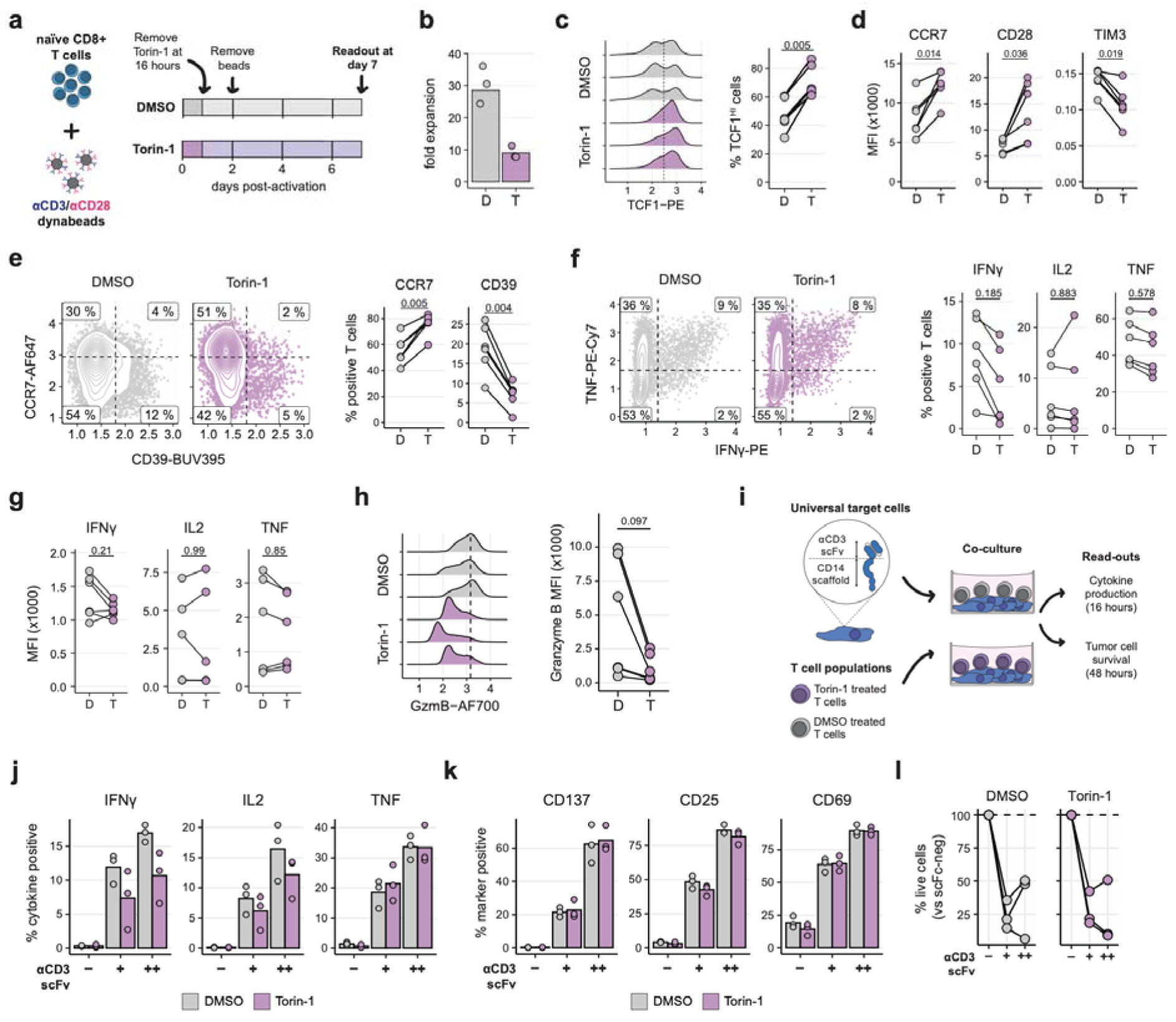
Transient mTOR inhibition during T cell priming imprints T cell state. (**a**) Experimental setup. Naïve CD8^+^ T cells were activated in the presence of Torin-1 or DMSO. After 16 hours Torin-1 and DMSO was removed, read-outs were performed at day 7 post-activation. (**b**) Fold-expansion of T cell cultures between day 0-7. (**c**) TCF1 protein expression, measured by intracellular flow-cytometry. Representative histograms of TCF1 protein staining (left) and percentage of TCF1-positive cells (right) are shown. Dashed line indicates the threshold for positive cells. (**d**) Surface staining for indicated proteins, measured by flow-cytometry. Median fluorescence intensity (MFI) of indicated proteins is shown. Solid lines connect individual donors. (**e**) Surface staining for indicated proteins, measured by flow-cytometry. Scatterplots with 2D kernel density contours for representative samples (left) and summarized percentages (right) are shown. Dashed lines indicate threshold for positive cells and solid lines connect individual donors. (**f**) Intracellular staining for indicated cytokines after a 3-hour re-stimulation within anti-CD3/CD28 antibodies in the presence of brefeldin A. Scatterplots with 2D kernel density contours for representative samples (left) and summarized percentages (right) are shown. (**g**) Median fluorescence intensity of cytokine positive populations shown in (f). (**h**) Granzyme B expression 7 days post-activation. Representative histograms (left) and summarized MFI (right) are shown. (**i**) Experimental setup for co-culture assay of T cells with mel888 melanoma cells that express a membrane tethered single-domain antibody directed against CD3. (**j**) Intracellular cytokine staining after a 16-hour co-culture in the presence of brefeldin A (added 15 minutes after start co-culture). Dots indicate individual donors. (**k**) Percentage positive T cells for indicated surface markers of activation after a 48-hour co-culture. Dots indicate individual donors. (**l**) Percentage of viable mel888 cells after a 48-hour co-culture. Solid lines connect individual donors, and dashed lines indicate 100% survival relative to wild type mel888 cells. Displayed data is compiled from of 2 independent experiments comprising 3-6 donors. P values indicated in (c-h) were calculated using a two-tailed paired Student’s t-test followed by Benjamini-Hochberg correction.

## DISCUSSION

In this study, we present a time-resolved dissection of matched transcriptome and proteome dynamics in activated human T cells. Our data show that TCR stimulation induces a highly ordered remodeling of the protein networks and places mTOR activity as a master regulator of T cell differentiation during early activation. Specifically, mTOR directs the choice between effector and multipotent T cell states within 24 hours of TCR stimulation by promoting translation (transcripts containing TOP motifs), proliferation (maintenance of SKP2), and differentiation (nuclear exclusion of FOXO1). These findings highlight that proteome remodeling during early T cell activation proceeds through tightly regulated molecular stages that are directly orchestrated through mTOR.

Our quantification of transcript–protein relationships during the first 24 hours of T cell activation uncovered that mRNA and protein abundance are often decoupled, with the majority of genes presenting limited concordance between mRNA and protein abundance. Interestingly, our analyses indicate that delayed protein expression accounts for a substantial fraction of the observed mRNA–protein associations. Nevertheless, even after accounting for this delay, approximately 50% of all differentially expressed proteins show no association with their corresponding mRNA levels. The most notable exception to this finding is cell-cycle–associated proteins, which display a strong positive correlation between mRNA and protein abundance, suggesting that their protein abundance is mainly driven by transcriptional control. Additionally, and in line with previous studies in static T cell states^10^, our temporally resolved data identify gene class-specific mRNA and protein expression patterns under dynamic conditions, strongly suggesting a dominant contribution of post-transcriptional and translational regulation. The molecular mechanisms governing these levels of regulation, although emerging approaches are now allowing their systematic investigation^49^. Irrespective of the regulatory pathways are yet to be defined, our study provides a valuable resource to study the kinetics of mRNA and protein expression during early T cell activation.

mTOR inhibition during T cell activation modulates the abundance of specific groups of proteins in a target-specific manner. For example, mTOR inhibition in murine T cells increased the expression of the CDK inhibitor p27, thereby inhibiting the cell cycle by altering the ratio of p27 and cyclin proteins^8^. Notably, we identify the ubiquitin ligase SKP2 as a likely effector linking mTOR signaling to cell-cycle-related protein expression during T cell activation. SKP2 has been described as a direct target of mTOR in gastric cancer, where mTOR activity promotes p27 degradation via SKP2 upregulation^37^. In agreement with this mechanism, Torin-1 treatment suppresses SKP2 accumulation and increased expression of p27 during T cell activation, leading to an impairment of the G1/S transition. Provided the marked reduction of SKP2 protein abundance following mTOR inhibition, it is tempting to speculate that mTOR directly stabilizes SKP2 through phosphorylation, as was reported for cancer cells^37^. Overall, our data point to SKP2 as a putative mediator through which mTOR coordinates activation-induced cell-cycle progression in human T cells.

Our findings support a staged temporal model of early T cell activation in which: (i) cell-cycle occurs rapidly (minutes - 2 hours), followed by (ii) translational capacity build-up (2-6 hours). Similarly to cell cycle genes, mTOR inhibition selectively represses protein modules involved in translation, including TOP-motif-containing transcripts^8,25^. Yet, proteins induced by T cell activation were largely unaffected by Torin-1 treatment, indicating that the enhanced translation of these proteins by TCR signaling pathways does not rely on mTOR activity. This is consistent with literature showing that NFAT and AP-1 activity is largely mTOR-independent^50,51^. Lastly, we show that mTOR (iii) instructs the commitment to differentiation at later stages of T cell activation (6-24 hours). At this time point, mTOR inhibition promotes the expression of the transcription factor TCF1, which mirrors the transcriptional and functional profile of memory-precursor cells observed in long-term rapamycin-treated murine models^21,22,27,52^. Mechanistically, we present that this shift of phenotype towards memory-like cells aligns with the nuclear translocation of FOXO1, which is prevented by mTORC2-mediated phosphorylation of AKT^22^. Building on this observation, we demonstrate that transient mTOR inhibition during T cell priming, limited to 16-hour treatment, is sufficient to durably skew T cell fate toward a phenotype resembling long-lived memory-precursor cells, while retaining their cytotoxic capacity. Thus, transient mTOR inhibition enhances T cell multipotency without compromising functional potency. Previous pre-clinical and clinical studies^32,43–45^ have shown that mTOR modulation during vaccination strategies enhances the frequency and longevity of memory-like T cells. However, the broad immunosuppressive effects associated with continuous mTOR inhibition strongly reduce the applicability of such drugs in a clinical setting. Our findings provide a rationale to overcome this limitation, proposing that short-term co-administration of mTOR inhibition during vaccination strategies should be sufficient to enhance the durability of vaccine-induced immunity.

## METHODS

### Cell culture

Peripheral blood mononuclear cells (PBMCs) from anonymized healthy donors were used in accordance with the Declaration of Helsinki (Seventh Revision, 2013) after written informed consent (Sanquin). PBMCs were isolated through Lymphoprep density gradient separation (Stemcell Technologies). For matched mass spectrometry and RNA-sequencing analysis, PMBCs from 3 pools of 40 donors were used. For flow cytometry analysis of Teff, PBMCs were pooled from 5 donors. Cells were used after cryopreservation. T cells were activated in 24-well plates were pre-coated overnight at 4°C with 2 µg/mL rat a-mouse IgG2a (MW1483, Sanquin) in phosphate-buffered saline (PBS). Plates were washed with PBS and coated for >3 h with 1 µg/mL αCD3 (HIT3a, Biolegend) at 37°C. 1.3×10^6^ PBMCs/well were seeded with 1 µg/mL soluble αCD28 (CD28.2, Biolegend) in 1 mL culture medium (Iscove’s Modified Dubecco’s Medium, IMDM) supplemented with 10% fetal bovine serum (FBS), 100 U/mL penicillin, 100 µg/mL streptomycin, and 2 mM L-glutamine. After 48h of activation at 37°C, 5% CO2, cells were harvested and cultured in standing T25/T75 tissue culture flasks (Thermo Scientific) at a density of 0.8×10^6^/mL in culture medium supplemented with 100 IU/mL recombinant human (rh)IL2 (Protech) and 10 ng/mL rhIL-15 (Protech). Culture media was refreshed every 2-3 days. Teff cells were cultured for 9 days without TCR triggering before re-activation and time-course sampling.

Naïve CD8^+^ T cells were isolated from PBMCs of individual donors using the BD IMag™ Human Naïve CD8+ T Cell Enrichment Set (BD Biosciences) following the manufacturers protocol. Cells were cultured in RPMI (Gibco) supplemented with 10% FBS, 100 U/mL penicillin, 100 µg/mL streptomycin, 50 IU/ml rhIL2 (Protech), 5 ng/ml rhIL15 (Gibco) and 10 ng/ml IL7 (Gibco). Isolated cells were activated using Dynabeads™ Human T-Activator CD3/CD28 for T Cell Expansion and Activation (Gibco) at a 1:1 T-cell-to-bead ratio, in the presence of 250 nM Torin-1 (dissolved in DMSO) or an equal v/v percentage of DMSO (0.05%). Torin-1 and DMSO was removed from cell cultures at the indicated timepoints by 3 sequential washes in complete culture media. Next, cells were pelleted and re-seeded in culture medium. CD8^+^ T cell cultures were passaged every 2-3 days and were kept at a density of 0.4-1.0×10^6^ cells/mL.

### T cell activation and time course sampling

Teff cells were restimulated with 1 μg/ml soluble anti-CD3ε (Pelicluster CD3, Sanquin) and 1 μg/mL anti-CD28 (clone 28.2 biolegend) for the indicated durations. For mTOR inhibition, 250nM Torin-1, or an equal v/v percentage of DMSO (0.05%), was added. For intracellular protein measurements, 1 μg/mL brefeldin A (BD Bioscience) and 1 μg/mL Monensin (Invitrogen) were added during the last two hours of activation. CD107a expression was measured by adding anti-CD107a (H4B3, BD Bioscience) from the start of activation. To measure translation efficiency, T cells were incubated with 5 μg/ml Puromycin dihydrochloride (Sigma) in culture media for 10 min at 37°C.

### RNA sequencing

1 x10^6^ T cells were used for RNA sequencing (Genewiz; Azenta Life Sciences). During RNA isolation and library preparation, ribosomal RNA (rRNA) was depleted. Approximately 30×10^6^ reads were obtained per sample. Sequencing adaptor trimming and low-quality read removal was performed using fastp^53^ (default settings). As rRNA depletion is not absolute, obtained reads were first mapped to a custom rRNA library (deposited to Zenodo) using STAR^54^. Unmapped reads were subsequently aligned to the GRCh38 primary assembly using the gencode.v47 basic annotation (both files downloaded from https://www.gencodegenes.org/human). Aligned reads were subsequently quantified using htseq-count^55^. Differential gene-expression analysis between indicated samples was performed using the R package limma^56^.

### Mass spectrometry data acquisition

For mass spectrometry, samples were lysed in 1% sodium deoxycholate (Bioworld), 10 mM TCEP (Thermo Scientific), 40 mM chloroacetamide (Sigma-Aldrich), 100 mM Tris-HCl pH 8.0 (Gibco). Lysates were incubated for 5Lminutes at 95 °C, sonicated for 10Lminutes in a sonifier bath (Branson model 2510), and digested overnight with 250 ng Trypsin/Lys-C (Promega) at 25 °C. Tryptic digests were desalted on an Assaymap BRAVO (Agilent) using C18 cartridges (5ul, Agilent) according to manufactures instructions. Concentrations were determined with a Pierce Quantitative Fluorometric Peptide Assay (Thermo). Peptides (500 ng) were loaded on Evotip Pure (Evosep) tips according to manufacturer’s guidelines and separated on a Performance Column (EV1115, EvoSep) using the 60 samples per day gradient. Buffer A consisted of 0.1% formic acid, buffer B of 0.1% formic acid in acetonitrile (Biosolve). Data was acquired on a timsTOF HT mass spectrometer (Bruker Daltonics) operated in diaPASEF mode, using a MS1 scan range of 100-1700 m/z. MS2 acquisition was performed using 32 pyDIAID-optimized^57^ mass and ion mobility windows, ranging from 400.2 to 1500.8 m/z and 0.70 to 1.50 1/k0 with a cycle time of 1.80 seconds. A collision energy of 20.00 eV at 0.6 1/k0 and 59 eV at 1.60 1/k0 was used.

### Mass spectrometry analysis

Raw mass spectrometry files were processed with DIA-NN 1.8.1, using the reviewed human proteome database (Swiss-ProtDatabase, 20,370 entries, release 2024.04.22). Standard settings and a generated library-based spectra search were used. Label free quantification values were log2 transformed. Missing values were imputed by a normal distribution (width=0.3, shift = 1.8), assuming these proteins were close to the detection limit. Differential abundance of proteins was tested using the R package limma^56^.

### Protein expression clusters and correlation to mRNA abundance

Proteins exhibiting differential expression in any comparison between time-points were selected. Cutoffs for differential expression were set at a Benjamini-Hochberg adjusted P < 0.05 and an absolute fold-change > 1.27, resulting in 2,650 proteins for further analysis. Protein expression was subsequently scaled across samples and divided into 5 clusters using kmeans clustering. Upon visual inspection of the expression profiles, clusters were defined as “early up", "gradual up", "late up", "gradual down" and "late down". Full list of classification in **Supplementary Table 3**. Enriched Reactome pathways in clusters were tested with the compareCluster function from the ClusterProlifer R package.

The within-gene correlation between protein and mRNA abundance was calculated as Spearman’s rank correlation coefficient. To account for the potentially delayed protein expression, ‘delayed correlations’ were calculated by shifting the time points obtained by RNA sequencing by 1 or 2 positions. This effectively tests if mRNA abundance is predictive of protein levels at later timepoints. Proteins and their relation to mRNA abundance were subsequently classified as “Positive association" (r > 0.5), "Delayed positive association" (r > 0.5 at time-point shifts of 1 or 2), "Negative association" (r < −0.5), "No association" (r > −0.5 and r < 0.5). Full list of classification in **Supplementary Table 9**.

### XGBoost modelling of protein expression clusters

To test the effects of Torin-1 treatment on the protein expression clusters, an XGBoost model was trained to classify cluster identity, using the protein abundance of the 2,650 differentially expressed proteins from Figure 1b. Before model training abundance values were scaled across samples. Model training was performed using the caret^58^ and xgboost^35^ R packages. Repeated cross-validation was applied during training (10 cross-validations, 2 repeats). Settings for the XGBoost model were: number of rounds = 1000, maximum tree depth = 1, subsample ratio of columns = 0.5, learning rate = 0.3, minimum loss reduction = 1, minimum sum of instance weight = 0.9, subsample ratio of the training instances = 1. Training objective was set to "multi:softprob" and the target metric was “Accuracy”. The resulting model had an accuracy of 0.901 and a kappa of 0.870. The obtained XGBoost model was subsequently used to predict class identity in the LC-MS data obtained in Figure 2 using the predict function implemented in the caret R package.

### Immunoblotting

Cell lysates (1×10^6^ cells/sample) were prepared by standard procedures using RIPA lysis buffer (Thermo) supplemented with protease and phosphatase inhibitors (Thermo). Lysates were run on 4–12% SDS-PAGE (Thermo). SDS-PAGE gels were directly transferred onto nitrocellulose membranes (iBlot2, Thermo). Membranes were blocked with 5% BSA TBST solution (Fraction V, Sigma). And incubated with α-SKP2 (Proteintech), or α-RhoGDI (MAB9959, Abnova), followed by α-Rabbit (4050-05, Southern Biotech) HRP-conjugated secondary antibodies.

### Flow cytometry and intracellular staining

T cells were washed with FACS buffer (PBS with 1% FBS and 2 mM EDTA) and labeled for 20 minutes at 4°C with α-CD4 (SK3, BD Horizon), α-CD8 (SK1, BD Horizon), α-CD69 (FN50, BD Horizon), α-TIM3 (73D, BD Horizon), α-CD137(4B4-1, BioLegend), α-CD27 (O323, Biolegend), α-CCR7 (3D12, BD Pharmigen), α-CD28 (CD28.2, BioLegend), α-CD39 (AI, BD Horizon), α-CD25 (BC96, BioLegend). Dead cells were excluded with Near-IR (Life Technologies). For intracellular staining, cells were fixed and permeabilized with Cytofix/Cytoperm kit (BD Biosciences) and stained with α-IFN-γ (4S.B3, BD Bioscience), α-TNF (MAb11, BD Bioscience), α-IL2 (MQ1-17H12, Biolegend) α-Puromycin (12D10, Merk). For transcription factor expression, cells were fixed and permeabilized with eBioscience™ Foxp3/ Transcription Factor Staining Buffer Set (Invitrogen) prior to staining with α-TCF1 (C63D9, Santa Cruz Biotechnology), and α-Tbet (4B10, Biolegend). For phospho-flow cytometry, cells were fixed with IC Fixation Buffer (eBioscience) for 10 minutes at 37°C and permeabilized with 90% ice-cold methanol for 30 minutes at 4°C. Cells were stained with α-4E-BP1 (V3NTY24, eBioscience) and α-AKT (SDRNR, eBioscience) for 1 hour at room temperature. For cell cycle staining, cells were fixed and permeabilized with eBioscience™ Foxp3/ Transcription Factor Staining Buffer Set (Invitrogen) prior to staining with α-Ki67(B56, BD Bioscience). Cells were then treated with 100 μg/ml of RNAse for 10 minutes at room temperature. Prior to acquisition, cells were stained with Propidium Iodide (Invitrogen). Acquisition was performed using BD LSRFortessa Cell Analyzer (BD Bioscience) or FACS Symphony A5 Cell Analyzer (BD Bioscience). Data were analyzed with FlowJo (BD Biosciences, version 10.8.1).

For quantification of FOXO1 nuclear localization, cells were labelled for 20 minutes at 4°C with RY586 mouse α-human CD8^+^ (568115, BD Biosciences). Cells were fixed and permeabilized using the eBioscience™ Foxp3/Transcription Factor Staining Buffer Set (Invitrogen) prior to staining for 1 hour at RT with α-FOXO1 (C29H4, Santa Cruz Biotechnology). Prior to acquisition, cells were stained with DAPI (62248, Thermo Scientific). Acquisition was performed using ImageStream MK II (Cytek Amnis). Data were analyzed with IDEAS (Amnis/ Luminex).

### Co-culture experiments

For co-culture experiments, a modified Mel888 line was used expressing a membrane-tethered anti-CD3 single-chain variable fragment (scFv) of the OKT3 clone, which was generated as previously described^59^, following the approach of Leitner et al^60^. 15,000 Mel888 cells were seeded per well on a flat-bottom 96-well tissue-culture plate. After 4 hours, 15,000 T cells were added to each well, resulting in a 1:1 effector:target ratio. T cells were seeded in the presence of 50 U/ml rhIL2. Plates were centrifuged at 10 × g for 1 minute (without brake) and incubated at 37C for 30 minutes. Next, Brefeldin A (BD Biosciences) was added to the culture, following manufacturers protocol. The co-culture was incubated at 37 °C for 16 hours. Co-cultures were harvested, and cytokine production was assessed using the BD Cytofix/Cytoperm™ Fixation/Permeabilization Kit (BD Biosciences). For measurements of T cell activation and melanoma cell killing, co-cultures were incubated for 48 hours before analysis. For these analyses, co-cultures were harvested and stained for the indicated activation markers. Before flow cytometry analysis, 5,000 Precision Count Beads™ (Biolegend) were added to allow for the calculation of surviving melanoma cells.

### Data Visualization and statistical analysis

Statistical analysis was performed using the rstatix R package, using two-tailed paired Student’s t-tests followed by Benjamini-Hochberg correction. P values are indicated in the figures. Data were visualized with ggplot2^61^ (version 3.4.2). Heatmaps were generated using the ComplexHeatmap R package.

## RESOURCE AVAILABILITY

### LEAD CONTACT

Information and requests for resources and reagents should be directed to and will be fulfilled by the lead contacts Monika C Wolkers and Kaspar Bresser

### MATERIALS AVAILABILITY

This study did not generate new unique reagents.

### DATA CODE AND AVAILABILITY

- The MS data have been deposited to the ProteomeXchange Consortium via the PRIDE^62^ partner repository with the dataset identifier PXD073124
- The RNA sequencing data have been deposited to DANS Data Station Life Sciences and are available at https://doi.org/10.17026/LS/PAYFTR
- All codes used in the analyses described in this study are available at https://github.com/kasbress/Tcell_Activation_mTOR
- Any additional information required to reanalyze the data reported in this paper is available from the lead contact upon request

## ACKNOWLEDGMENTS

We thank Erik Mul from the Sanquin Central Facility for analysis of the FOXO1 nuclear localization data. This research was supported by Oncode Institute, the European Research Council consolidator award PRINTERS 817533, and Landsteiner Foundation for Blood Transfusion (LSBR) Research grant 2202 to M.C.W.; and the Cancer Center Amsterdam (CCA) research grant CCA2023-9-91 to K.B.

## AUTHOR CONTRIBUTIONS

Conceptualization, A.P.J, M.V.L., K.B., M.C.W.; Methodology, A.P.J, M.V.L., K.B., and M.C.W.; Investigation, A.P.J, M.V.L., L.W., K.B., C.v.d.Z., A.J.H.; Formal Analysis, A.P.J, M.V.L., K.B., A.J.H.; Validation, A.P.J, M.V.L., K.B..; Writing-Original Draft, A.P.J, M.V.L., K.B. Writing – Review & Editing, A.P.J, M.V.L., K.B. and M.C.W.; Supervision, K.B., and M.C.W.; Funding Acquisition, M.C.W and K.B..

## DECLARATION OF INTEREST

The authors declare no competing interests.

---

**Supplementary Figure 1.**
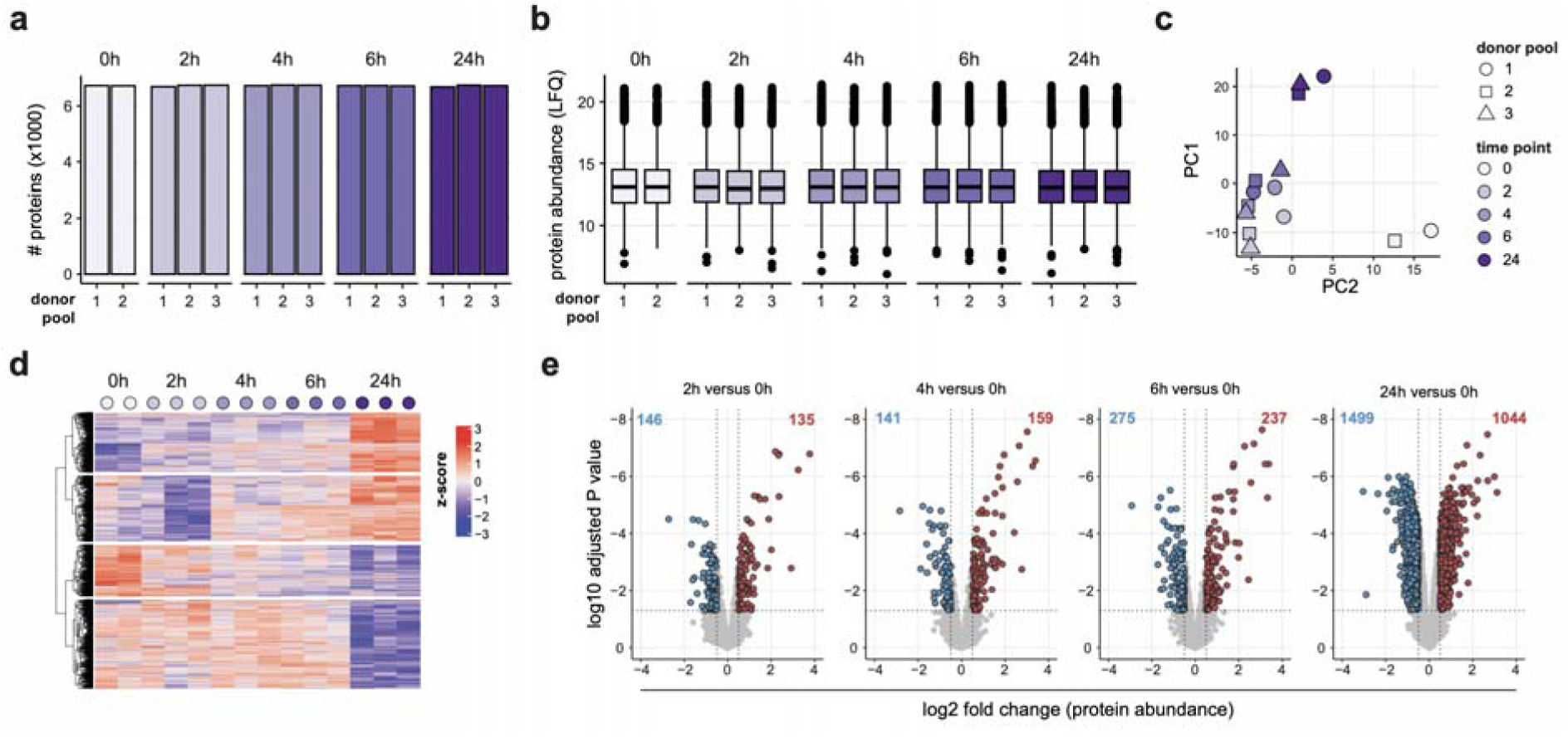
LC-MS dataset of T cell activation time course. (**a-b**) Number of identified proteins (a) and intensity value (LFQ) (b) at indicated activation time points of CD3/CD28-activated T cells from LC-MS data. (**c**) Principal component analysis of LC-MS data. Treatment conditions are indicated by color, donor pools (experimental replicates) by shapes. (**d**) Clustering analysis of all differentially expressed proteins from the LC-MS data. (**e**) Differentially expressed proteins of LC-MS data, comparing each activation time point (2, 4, 6, and 24 hours) to the 0-hour time point. Colored numbers indicate the number of proteins that pass the significance threshold (adjusted P < 0.05).

**Supplementary Figure 2.**
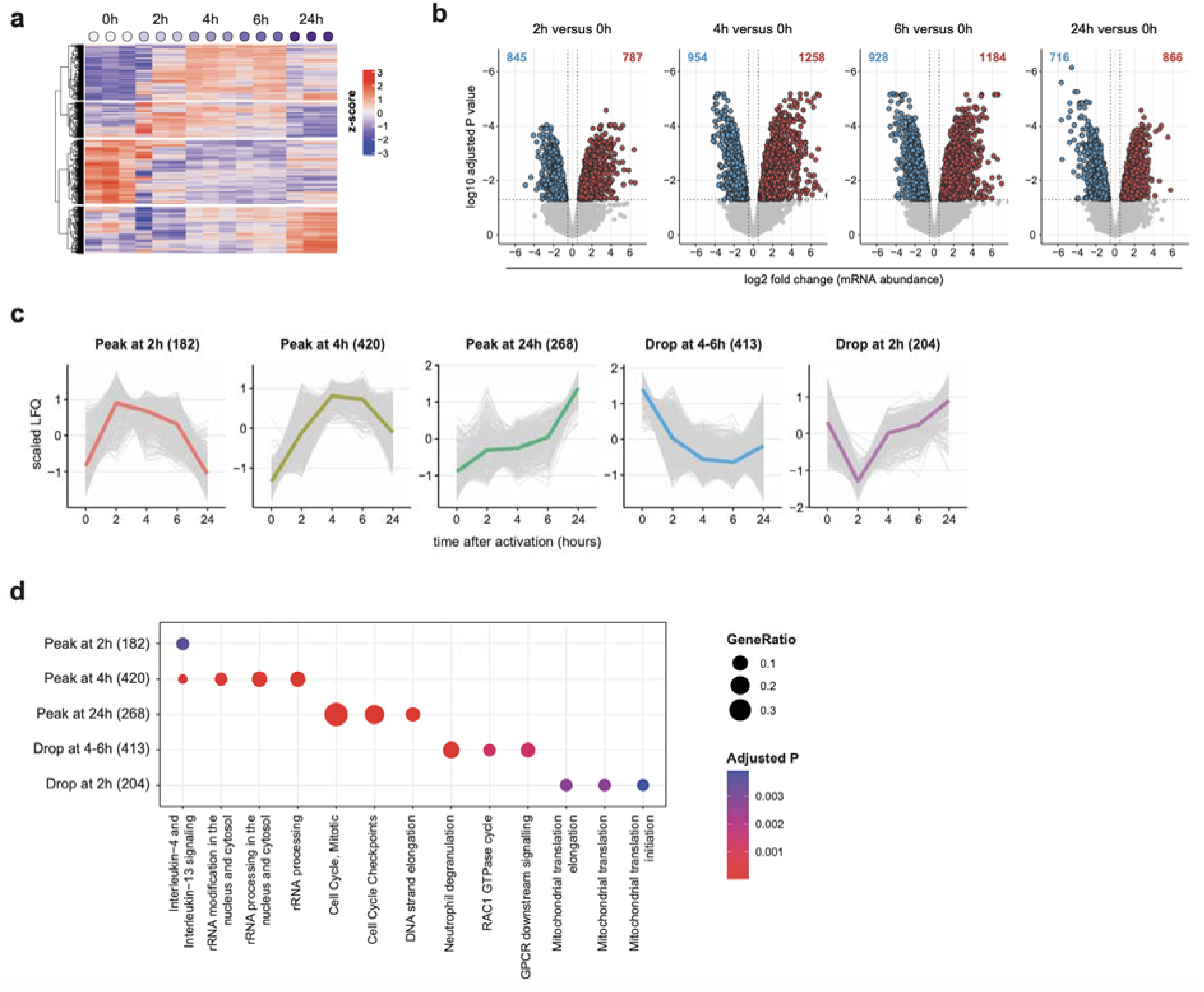
mRNA-seq dataset of T cell activation time-course. (**a**) Clustering analysis of all differentially expressed genes of activation time points (0, 2, 3, 4, 6 and 24 hours) from the RNA-seq data. (**b**) Differentially expressed genes of RNA-seq data, comparing the activated time points (2, 4, 6, and 24 hours) to the 0-hour time point. Colored numbers indicate the number of transcripts that pass the significance threshold (adjusted P < 0.05). (**c**) Differentially expressed transcripts (n = 1,487; FDR < 0.05) were clustered into five temporal expression modules using hierarchical clustering. See **Supplementary Table 7** for cluster annotation. Grey lines indicate individual transcripts; colored lines indicate medians. (**d**) Enrichment of Reactome pathways in each module. A maximum of 4 pathways is shown for each cluster. See **Supplementary Table 8** for full list.

**Supplementary Figure 3.**
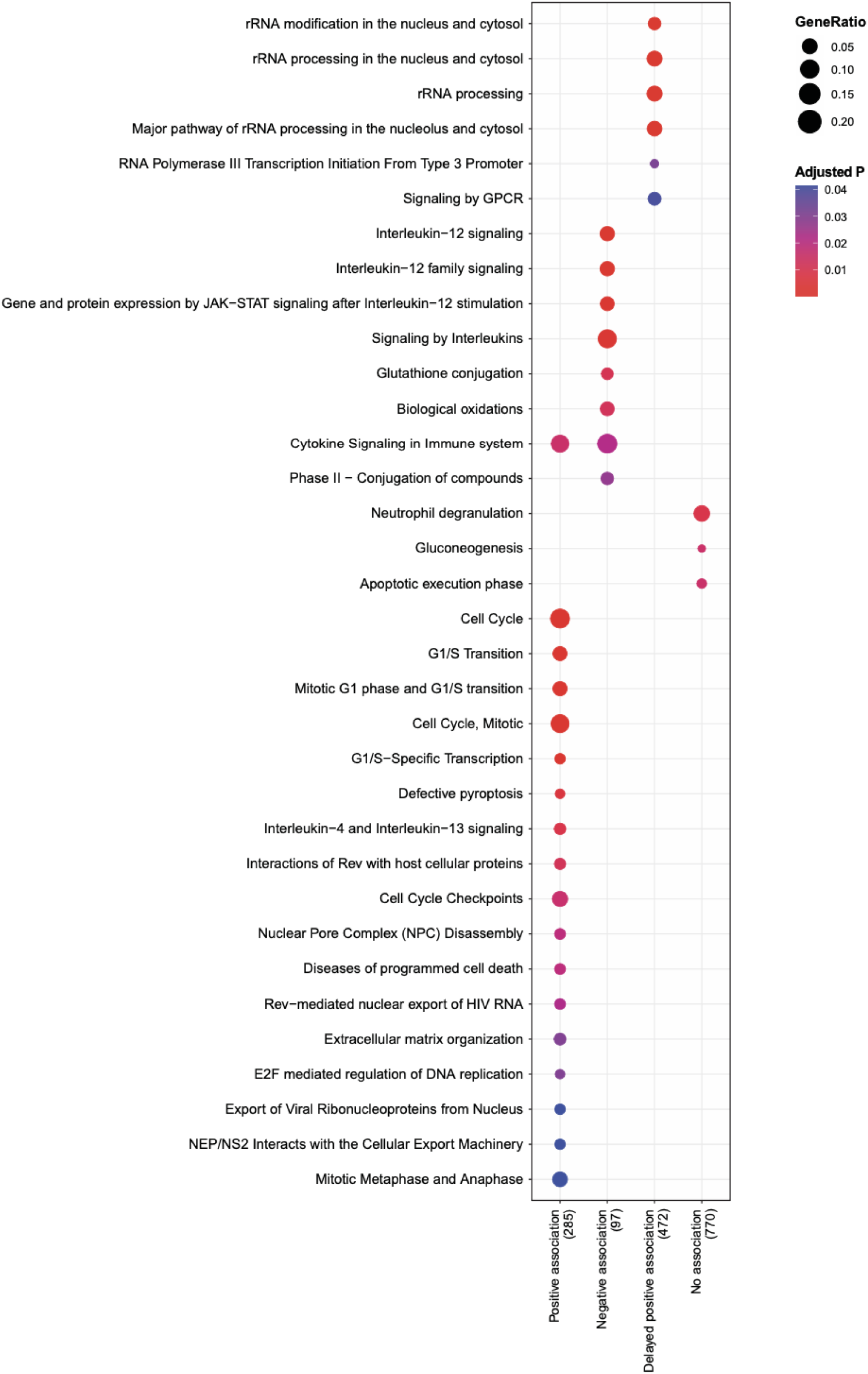
Pathway analysis of mRNA-protein association clusters. Enrichment of Reactome pathways in each cluster defined in Figure 1h-i. A maximum of 15 pathways is shown for each cluster. See **Supplementary Table 9** for full list.

**Supplementary Figure 4.**
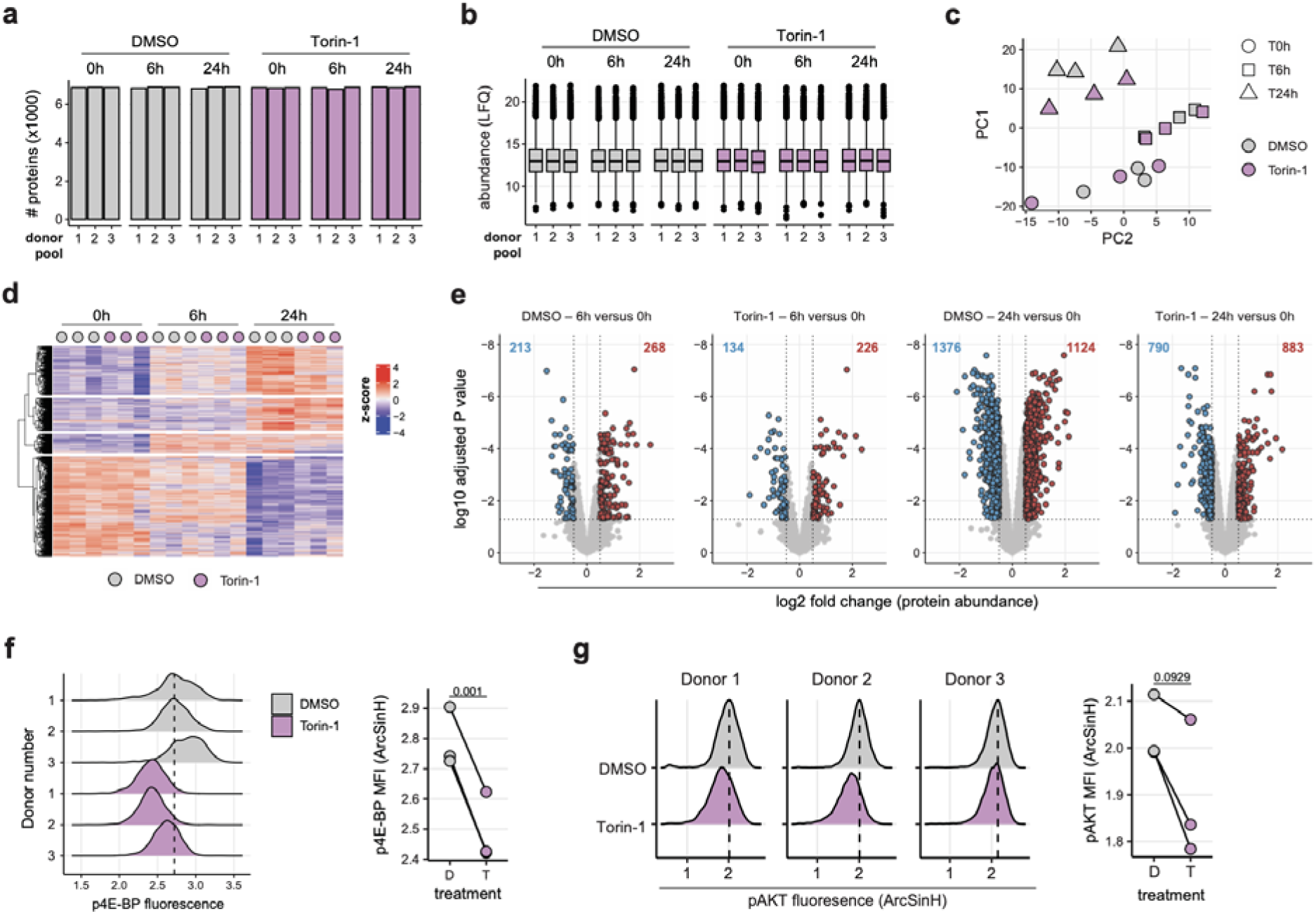
LC-MS dataset of T cell activation with mTOR inhibition. (**a-b**) Number of identified proteins (a) and corresponding LFQ intensity value (b) of Torin-1 treated T cells, at indicated time points from LC-MS data. (**c**) Principal component analysis of LC-MS data. Treatment conditions are indicated by color, time points by shapes. (**d**) Clustering analysis of all differentially expressed proteins from the LC-MS data, comparing Torin-1 treated (purple) with the DMSO control (grey) samples. (**e**) Differentially expressed proteins of LC-MS data, comparing activation time points (6 and 24 hours) to the 0-hour time point within each treatment condition. (**f-g**) Teff cells re-activated with anti-CD3/CD28 for 24 hours in the presence of Torin-1 or DMSO control. (**f**) Intracellular staining for p-4E-BP1 (Thr36/Thr45 phosphorylation site), measured by flow-cytometry. Representative histograms of anti-p-4E-BP1 fluorescence intensity (a) and median fluorescence intensity (b) are shown. Dashed lines indicate the maximum density of fluorescence for the DMSO controls and solid lines connect individual donors. (**g**) Intracellular staining of p-AKT (Ser473 phosphorylation site), measured by flow-cytometry. Representative histogram of anti-p-AKT fluorescence intensity (c) and median fluorescence intensity (d) are shown. Dashed lines indicate the maximum density of fluorescence for the DMSO controls and solid lines connect individual donors. Displayed data in (f-g) is representative of 2 independent experiment comprising 3 pools of 5 donors. P values indicated in (f-g) were calculated using a two-tailed paired Student’s t-test followed by Benjamini-Hochberg correction.

**Supplementary Figure 5.**
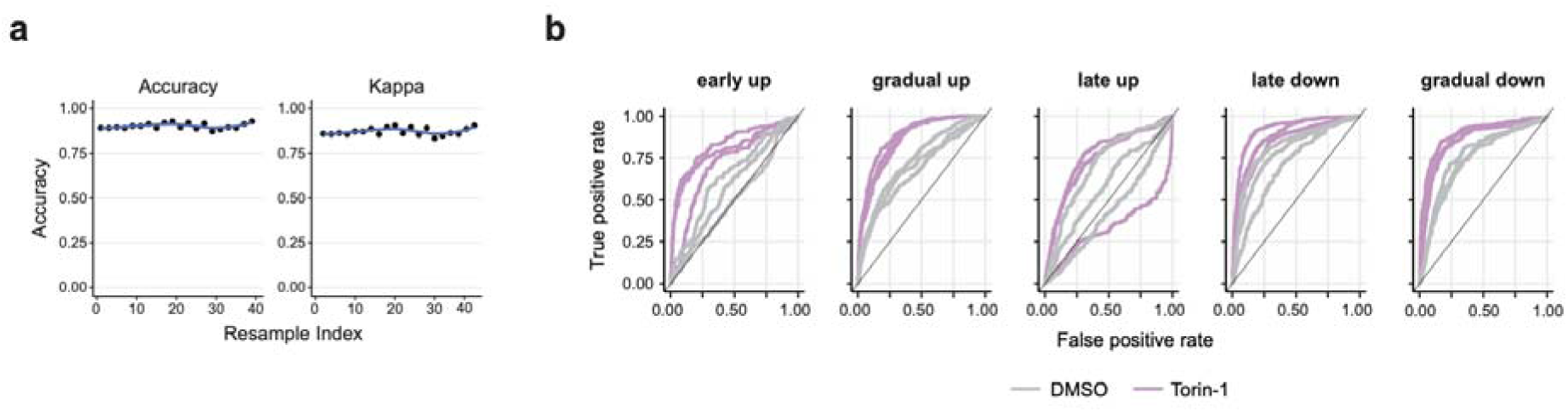
Modeling of protein expression modules using XGBoost. LC-MS data from Figure 1 were used to train a XGBoost classifier for the 5 indicated protein expression modules. This model was then used to classify the DMSO/Torin-1 LC-MS data (Figure 2). (**a**) XGBoost model performance metrics during resampling. (**b**) Model performance as receiver operating characteristic (ROC) curves. Diagonal back lines denote random predictions.

## Notes

### Competing Interest Statement

The authors have declared no competing interest.

